# Deficiency in DNA mismatch repair of methylation damage is a major mutational process in cancer

**DOI:** 10.1101/2020.11.18.388108

**Authors:** Hu Fang, Xiaoqiang Zhu, Jieun Oh, Jayne A. Barbour, Jason W. H. Wong

## Abstract

DNA mismatch repair (MMR) is essential for maintaining genome integrity with its deficiency predisposing to cancer^1^. MMR is well known for its role in the post-replicative repair of mismatched base pairs that escape proofreading by DNA polymerases following cell division^2^. Yet, cancer genome sequencing has revealed that MMR deficient cancers not only have high mutation burden but also harbour multiple mutational signatures^3^, suggesting that MMR has pleotropic effects on DNA repair. The mechanisms underlying these mutational signatures have remained unclear despite studies using a range of *in vitro*^4,5^ and *in vivo*^6^ models of MMR deficiency. Here, using mutation data from cancer genomes, we identify a previously unknown function of MMR, showing that the loss of non-canonical replication-independent MMR activity is a major mutational process in human cancers. MMR is comprised of the MutSα (MSH2/MSH6) and MutLα (MLH1/PMS2) complexes^7^. Cancers with deficiency of MutSα exhibit mutational signature contributions distinct from those deficient of MutLα. This disparity is attributed to mutations arising from the unrepaired deamination of 5-methylcytosine (5mC), i.e. methylation damage, as opposed to replicative errors by DNA polymerases induced mismatches. Repair of methylation damage is strongly associated with H3K36me3 chromatin but independent of binding of MBD4, a DNA glycosylase that recognise 5mC and can repair methylation damage. As H3K36me3 recruits MutSα, our results suggest that MutSα is the essential factor in mediating the repair of methylation damage. Cell line models of MMR deficiency display little evidence of 5mC deamination-induced mutations as their rapid rate of proliferation limits for the opportunity for methylation damage. We thus uncover a non-canonical role of MMR in the protection against methylation damage in non-dividing cells.

## Introduction

DNA mismatch repair (MMR) is a highly conserved DNA repair pathway, well known for its ability to recognise and remove errors from the newly synthesised DNA strand during replication^7^. The general mechanism of MMR in the correction of DNA replication errors in humans is well established. The process is initiated by the MutSα heterodimer, comprised of MSH2 and MSH6, which recognise mis-incorporated bases in double stranded DNA behind the replication fork. Subsequently, the MutLα heterodimer, which is comprised of MLH1 and PMS2, is recruited to excise the sequence surrounding the mutated base. The mismatched section of the daughter strand is digested by EXO1 exonuclease and the gap filled by a DNA polymerase^7^. MMR is also known to have non-canonical roles outside the context of DNA replication^8^. One of the better-known functions of non-canonical MMR (ncMMR) is facilitating somatic hyper-mutation of the immunoglobulin locus in lymphoid cells^9^. NcMMR has also been shown to be activated by DNA lesions resulting in an error-prone repair process that leads to the formation of A:T mutations^10–12^. While these studies have found that ncMMR is generally associated with increased mutagenesis through the recruitment of error-prone DNA polymerase eta (POLH), more recently, it has also emerged that ncMMR is capable of protecting actively transcribed genes by removing DNA lesions in a transcription dependent manner^13^. The mechanistic details of how ncMMR achieves error-free repair remains unclear, but there is also evidence that ncMMR can facilitate active DNA demethylation^14^, suggesting that there are multiple ncMMR pathways and a pathway for high-fidelity ncMMR may exist.

The use of somatic mutational signatures has contributed significantly to our understanding of the underlying mutational processes in cancer^15^. Currently seven single based substitution (SBS) mutational signatures have been associated with MMRd across various cancer types^3^. MMRd mutational signatures, SBS6 and SBS15 are characterised by high frequency of C>T mutations, SBS21 and SBS26 have a dominant mutation spectrum of T>C, while SBS44 has relatively high contributions from C>A, C>T and T>C mutations. SBS14 and SBS20 are associated with MMRd concurrent with polymerase epsilon (POLE) and polymerase delta (POLD1) exonuclease mutations, respectively. Recently, it was shown that two mutational processes largely underlie the MMRd specific mutational signatures^16^. Both of these mutational processes are believed to be associated with DNA polymerase associated replication errors, yet, only one has been reproducible *in vitro* using clonal models of MMRd cell lines^5,16^. Thus, the mechanisms underlying the mutational processes and the mutational signatures observed in human MMRd cancers remain unclear.

In this study, we provide evidence that ncMMR is required for the repair of 5-methylcytosine (5mC) deamination induced G:T mismatches (we will refer to this as 5mC deamination damage) outside of the context of DNA replication. We show that mutations arising from unrepaired 5mC deamination events are prevalent in MMRd cancers, particularly those deficient of MutSα. Therefore, the deficiency of ncMMR activity is a major mutational process in MMRd cancers.

## Results

### Profile of mismatch repair genes in MSI-H tumors

We first investigated three tumor types (CRC, colorectal cancer; STAD, stomach adenocarcinoma; UCEC, uterine corpus endometrial carcinoma) as they have been recognised as MSI prone and contain the majority of MSI events^17^. A set of 316 cancer samples with MSI-High (MSI-H) phenotype was obtained from TCGA Pan-cancer cohort for these tumor types. In order to focus on samples where the major mutational processes is MMRd, we excluded samples with concurrent POLE and POLD1 exonuclease domain mutations as defined by high contributions from SBS14 and SBS20, respectively^18^. The remaining samples were stratified into MutLα and MutSα mutants after careful review of the underlying mutations, RNA expression and DNA methylation status of the four canonical mismatch repair genes (*MLH1, PMS2, MSH2* and *MSH6*) (see Methods, **Extended Data Fig 1, Supp Table 1**). In the end, 197 MutLα and 18 MutSα deficient cancer samples were identified. In addition, 14 samples were found with both MutLα and MutSα defects and 37 samples were unable to be conclusively classified based on the available data.

As expected, all the MSI-H samples showed high mutation load for both indels and single base substitutions with high proportions of C>T and T>C **(Fig 1A)**. MLH1 promoter methylation (defined as beta value > 0.25) was observed in 78% (149/191) of MSI-H samples with available methylation data and this is in line with previous studies^19,20^. For the curated MutLα deficient cancers, 93.4% (184/197) had aberrant expression of *MLH1* while the majority with MutSα deficiency harboured truncating mutations in either MSH2 or MSH6 **(Fig 1A)**. In order to compare the mutation spectrum for MutLα and MutSα deficient samples, principal component analysis was performed based on trinucleotide context point mutation frequencies. Samples with MutSα deficiency were clustered together with a relatively high frequency of C>T mutations, while MutLα deficient cancers were more distributed with a broader mutation spectrum **(Fig 1B)**. These results suggest that there may be differences underlying the mutational process of MutLα and MutSα deficiency.

**Fig 1.**
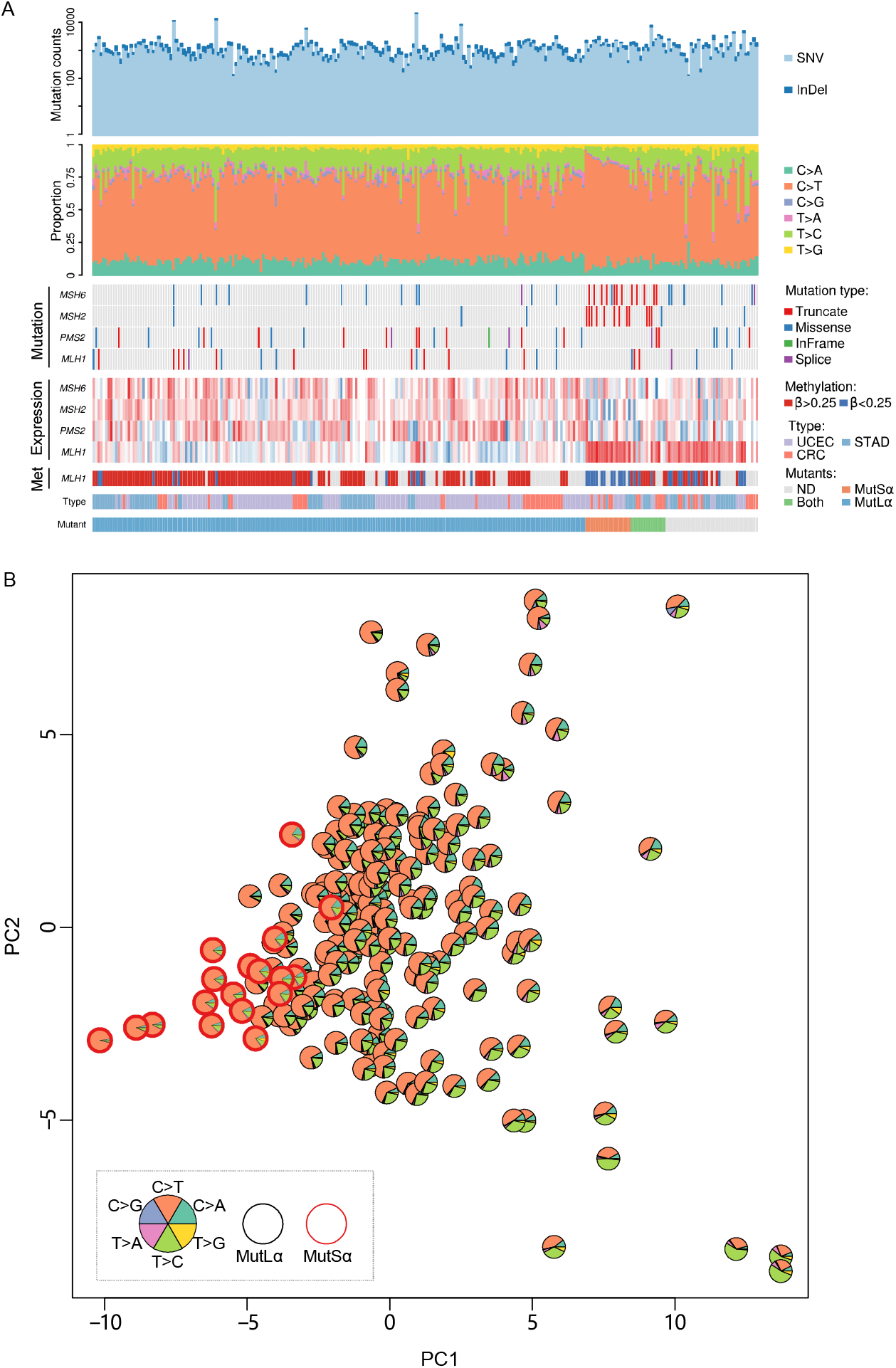
Landscape of MSI-H samples. (**A**) Profile of mutation burden and six types of mutation frequency, as well as the aberrant status of mismatch repair genes including DNA mutation, RNA expression and methylation. Cancer types and mutants classification are also indicated. (**B**) Principal component analysis of MSI-H cancer samples based on the frequency of 96 types of mutational contexts. The fractions of the six types of mutations are represented by the area of the sectors and MutSα mutants are highlighted in red.

### MutSα and MutLα deficient cancers display differential mutational signatures

To determine if MutSα and MutLα deficient cancers have different mutational processes, we used Sigfit^21^ to assign the somatic mutations of each sample to the five MMRd associated SBS mutational signatures along with the age-associated SBS1, which is present in most cancers. SBS1 contributed to a surprisingly high proportion of mutations in many MMR samples (**Fig 2A**), however, this may reflect difficulty in resolving SBS1 and SBS6 as the two signatures show a high degree of similarity. Generally, MutSα deficient cancers had the highest contribution from SBS1+SBS6 (**Fig 2A**). To simplify the representation of mutational processes in MMRd cancers, we adopted the use of two signatures proposed by Nemeth et al.^16^. Using the non-negative matrix factorization (NMF) algorithm, two *de novo* signatures were decomposed from the mutations of the MMRd samples. In line with Nemeth et al.^16^, two distinctive *de novo* signatures were obtained with signature A (SigA), characterised by a high frequency of C>T mutations, particularly in CpG sites while signature B (SigB) showed a broader spectrum with C>A, C>T and T>C mutations **(Fig 2B)**. We then calculated the cosine similarity between the reconstructed spectrum derived from these two signatures and the real spectrum for each sample genome. Most of the samples had high cosine similarity above 0.85 (**Extended Data Fig 2A**). Next, we compared the newly decomposed signatures with the previously reported MMRd related signatures^3^. SigA showed relatively high cosine similarity with SBS6 and SBS15 while SigB was more alike with SBS21 and SBS26 (**Extended Data Fig 2B**).

**Fig 2.**
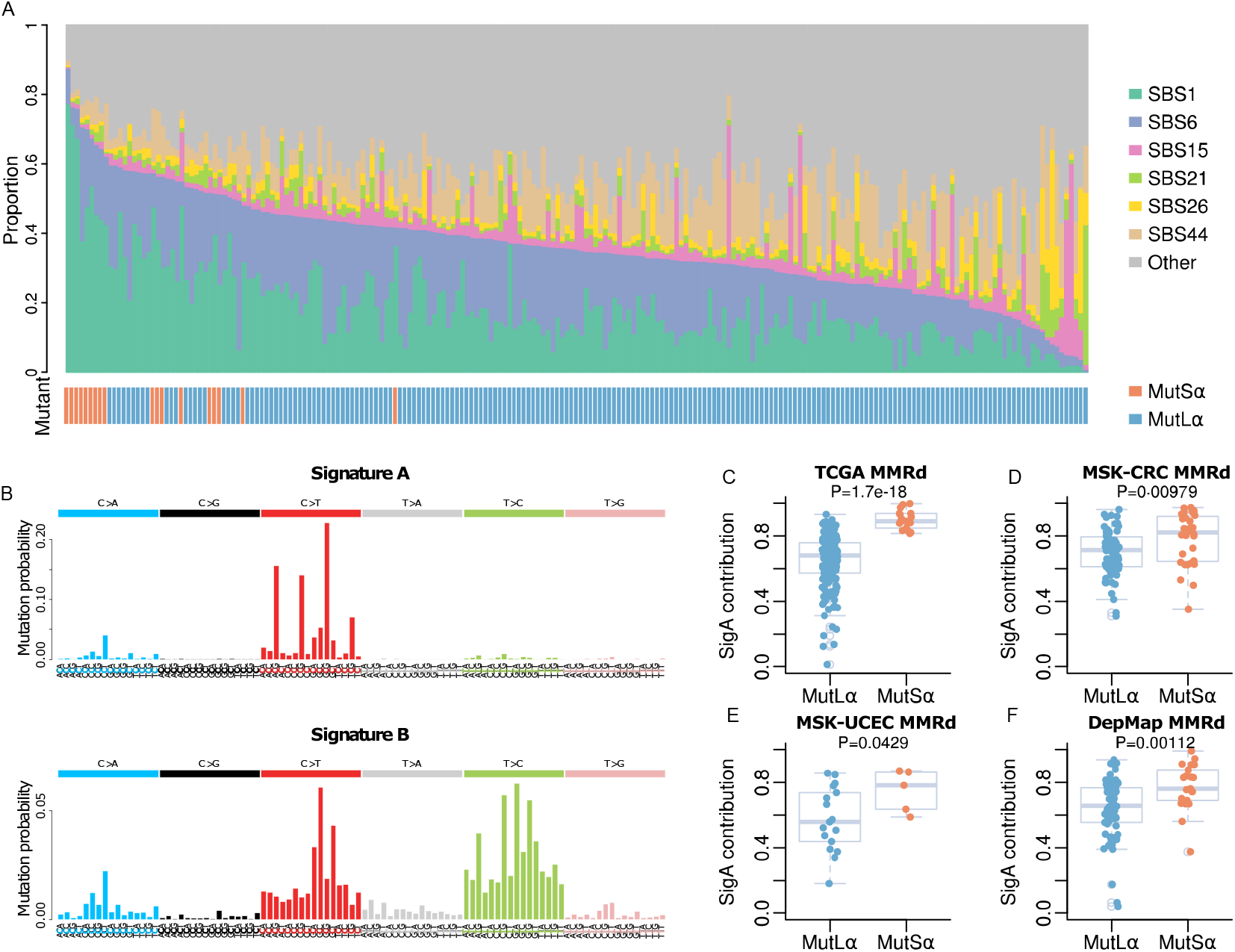
Mutation signature contribution in MutSα and MutLα mutants. (**A**) Fraction of MMRd associated signatures and age-related signature SBS1 contribution in MutSα and MutLα mutants. (**B**) The spectrum of *de novo* signatures extracted from TCGA MSI-H cancer samples. (**C-F**) The boxplot of SigA contribution for MutSα and MutLα mutants in TCGA-MMRd, MSK-CRC, MSK-UCEC and Depmap MMRd cohort. P-values were calculated by two-tailed Student’s t-test.

As we found that the mutation spectrum of MutSα deficient samples had generally higher proportion of C>T mutations and SBS1+SBS6 relative to MutLα deficient samples (**Fig 1A-B and Fig 2A)**, we sought to examine the contribution of the two *de novo* signatures in each MMR sample. Samples with MutSα deficiency had significantly higher contribution of SigA relative to those with MutLα deficiency (p < 0.001, Student’s t-test, **Fig 2C**), with samples deficient of both complexes having SigA contribution less than MutSα but not significantly different to MutLα deficient samples (p < 0.01 and p = 0.41, respectively, Student’s t-test, **Extended Data Fig 2C**). To exclude any potential cancer specific effect, we examined the signature contribution in CRC, STAD and UCEC separately, and found that SigA is significantly more enriched in MutSα compared with MutLα deficient samples across all cancer types (p < 0.01, Student’s t-test, **Extended Data Fig 2D-F**). We next expanded our data to three independent cohorts to validate this observation. Due to limited availability of data types for these cohorts, the approach to classify MutSα and MutLα deficiency status is slightly different from the TCGA dataset (see Methods). MSK-CRC^22^ and MSK-UCEC^23^ cohorts contain 99 CRC and 22 UCEC MSI-H samples, respectively, with only targeted sequencing data available. After fitting the mutations from the samples to SigA and SigB, both the cohorts showed significant enrichment of SigA in MutSα compared with MutLα deficient samples (p < 0.001 and p = 0.04 for CRC and UCEC respectively, Student’s t-test, **Fig 2D and E**). Furthermore, the analysis was also performed on the Depmap^24^ cohort comprising of 99 MMRd cell lines across 16 cancer types. Again, MutSα deficient samples had significant enrichment in SigA compared with MutLα deficient samples (p = 0.001, Student’s t-test, **Fig 2F**). Finally, as complex MSH2 and MSH6 mutations are a frequent mechanism of MSI in prostate cancer^25^, we identified a further 4 MutSα mutant samples in the TCGA prostate adenocarcinoma (PRAD) cohort and confirmed them all to have high SigA contribution (>0.845, **Extended Data Fig 2G**). Together, these results suggest that these two MMR associated mutational processes contribute to different extent to the overall mutation spectrum of MutSα and MutLα deficient cancers.

### CpG C>T mutations in MutSα mutants show no replication strand bias compared with non-CpG C>T mutations

Mutation density varies across cancer genomes^26^. Due to differential MMR efficiency, strong correlation is found between DNA replication timing and mutation density where late replicating regions have higher mutation density compared with early replicating regions of the genome^27^. In MMRd cancers, the association of mutation density and replication timing becomes less apparent for MMR dependent mutational processes. Given that we found different contributions of mutational processes in MutSα and MutLα deficient cancers, we sought to determine how the processes are influenced by replication timing. As expected, higher mutation density was observed in late replicating regions compared with early replicating regions in MMR proficient microsatellite stable (MSS) cancers but the difference was reduced in MutSα and MutLα deficient MSI samples (**Extended Data Fig 3A**). We observed significant difference in the dependence of the mutation load and replication time between MutSα and MutLα (**Extended Data Fig 3B**) and ascribe this difference to CpG C>T mutations but not non-CpG C>T or T>C mutations (**Extended Data Fig 3C-O**). These data suggest that there may be differences in the dependence of CpG C>T mutations on replication compared with other types of substitution mutations.

To further examine the relationship between mutation formation in MMRd cancers and DNA replication, we next investigated the replication strand bias of mutations. Mutations in MMRd samples have generally been attributed to unrepaired errors that have escaped from polymerase proofreading during DNA replication^28^. Due to the differential fidelity of polymerases during DNA synthesis, mutation load in the leading strand and lagging strand are expected to be asymmetric^29^. We examined the replication asymmetry of C>T/G>A and A>G/T>C mutations in the leading and lagging strands. Both mutation types showed strand bias with the leading strand, generating more C>T and A>G (p < 0.05 for both, Chi-squared test, **Fig 3A**). The lack of strand bias for exome-wide simulated mutations confirmed that this bias is related to replication rather than sequence composition (**Fig 3B**). We next compared the strand bias for each individual MutSα and MutLα deficient samples. While there was no difference in replication bias in A>G/T>C mutations between the MutLα and MutSα deficient samples (p = 0.849, Student’s t-test, **Fig 3C**), the bias of C>T/G>A mutation is significantly different, with MutSα showing less asymmetry than MutLα deficient samples (p=0.002, Student’s t-test, **Fig 3D**). Further, we compared CpG C>T and non-CpG C>T replication bias individually for MutSα and MutLα deficient samples. CpG C>T mutations showed significantly less bias compared with non-CpG C>T mutations (p<0.001 for both MutSα and MutLα, Student’s t-test, **Fig 3E**). We further observed significant CpG C>T strand bias difference between MutSα and MutLα deficient samples (P<0.001, Student’s t-test) while there was no significant difference for non-CpG C>T mutations (**Fig 3E)**. Although there was insufficient whole genome sequenced (WGS) samples with MutSα deficiency (n = 1), with increased number of mutations we were able to assess strand bias across all mutation types (**Extended Data Fig 4A-F**) and consistent results were observed in the strand bias difference between CpG C>T and non-CpG C>T mutations (MutLα, p < 0.001, Student’s t-test, **Extended Data Fig 4G)**. Reduced strand bias of CpG C>T mutations was also observed in WGS MSS samples (p < 0.001, **Extended Data Fig 4H**). Furthermore, as some MutLα deficient samples had relatively high SigA contribution, we directly correlated SigA contribution with the degree of CpG C>T replication bias. Negative correlation was observed with samples with high proportions of SigA showing less bias (R = −0.29, p < 0.0001, **Extended Data Fig 4I**). Together these results suggest that CpG C>T mutations associated with SigA in human MMRd cancers may not have arisen from unrepaired DNA polymerase errors as they do not exhibit the characteristic replication asymmetry found for other types of substitution mutations.

**Fig 3.**
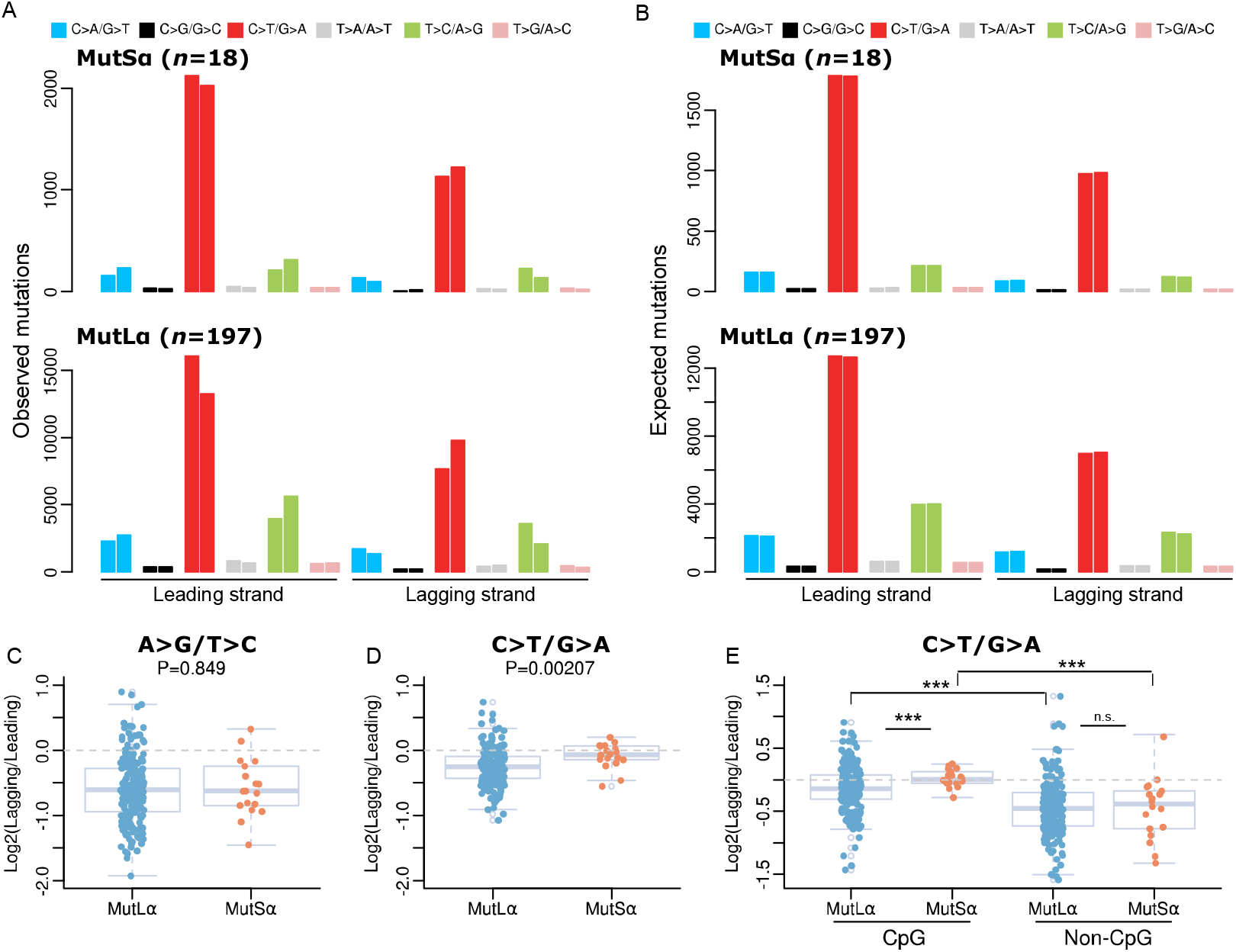
Replication asymmetry for MMR deficiency samples. Landscape of replication asymmetry for all observed mutations (**A**) and expected mutations (**B**) in MutSα and MutLα mutants. The expected mutations were obtained from simulation data that consider the abundance of tri-nucleotide mutational contexts. (**C-D**) Boxplot of replication stand bias for A>G/T>C and C>T/G>A mutations in MutSα and MutLα mutants. (**E**) Boxplot of replication stand bias for CpG C>T and non-CpG C>T mutations in MutSα and MutLα mutants. *** <0.001, n.s. >0.05, two-tailed Student’s t-test.

### Most CpG C>T mutations in MMRd cancers are caused by the deamination of 5-methylcytosine

5mC has the tendency to undergo spontaneous deamination into thymine and is a major mutagenic process in the human genome^30^. Methyl-binding domain 4 (MBD4) is a key DNA glycosylase responsible for the removal of 5mC deamination damage^31^. Cancer patients with biallelic germline MBD4 deficiency present with extremely high frequency of CpG C>T mutations, providing a model for mutations induced by 5mC deamination^32^. CpG C>T mutations in MBD4 deficient cancers are likely to arise outside the context of DNA replication thus replication asymmetry would not be expected. To test if this is the case, we obtained somatic mutations from whole genome sequenced MBD4-deficient cancers^32^ and compared the CpG C>T replication strand bias with MMRd mutants. We also included POLE exonuclease domain mutant cancers as their somatic mutations are known to be predominantly leading strand biased^33^ and it has been proposed that POLE mutants are particularly erroneous when replicating 5mC^34^. MBD4 mutants show little strand bias for all mutation types while there was substantial strand bias for POLE mutants (**Fig 4A**), an observation not present in simulated mutations (**Fig 4B**). The strand bias of CpG C>T mutations was close to zero for MBD4 mutants while, as expected, POLE mutants presented strong leading strand bias for both CpG and Non-CpG C>T mutations (**Fig 4C-D**). Although CpG C>T mutations for POLE mutants were highly enriched in TCG trinucleotide context, we observed consistent CpG C>T mutation strand bias in other contexts (**Extended Data Fig 5**). These results suggest that the pattern of MMRd CpG C>T mutations is more similar to MBD4 mutants where the mutations arise from the deamination of 5mC and are less dependent on process of DNA replication.

**Fig 4.**
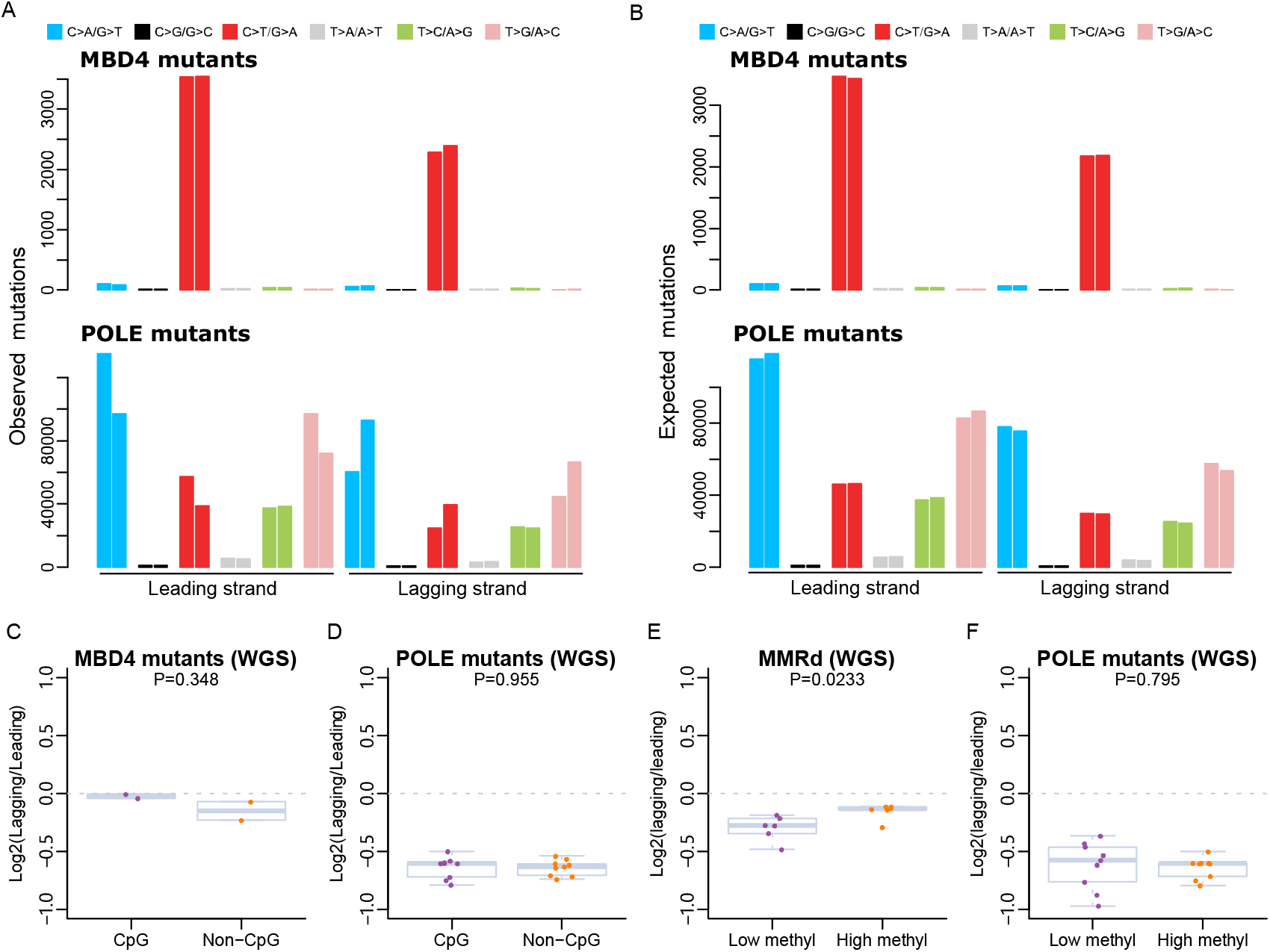
Replication asymmetry for MBD4 and POLE mutants. Landscape of replication asymmetry for all observed mutations (**A**) and expected mutations (**B**) in MBD4 and POLE mutants. The expected mutations are obtained from simulation data that consider the abundance of tri-nucleotide mutational contexts. (**C-D**) Boxplot of replication stand bias for CpG C>T and non-CpG C>T mutations in MBD4 and POLE mutants. (**E-F**) Boxplot of replication stand bias for CpG C>T mutations in highly methylated and lowly methylated regions for MMR deficiency samples and POLE mutants. Sites with beta value >0.3 are defined as highly methylated while <0.3 as lowly methylated. The range of mutation counts in lowly and highly methylated sites for calculating strand bias in MMRd samples were (52-156) and (2,347-6,145) respectively, and for POLE mutants (72-703) and (4,032-39,580) respectively.

As CpG C>T mutation rate increases with methylation level in MMRd samples^35^, we compared replication strand bias of CpG C>T mutations in lowly and highly methylated sites in MMRd and POLE mutant cancers. Mutations at highly methylated regions showed significantly less strand bias compared with lowly methylated sites for MMRd samples (p = 0.0233, Student’s t-test, **Fig 4E**) while POLE mutants presented comparably strong strand bias at both lowly and highly methylated regions (p = 0.80, Student’s t-test, **Fig 4F**). This further supports our hypothesis that CpG C>T mutations in MMR deficient cancers arise from replication independent deamination of 5mC.

### MMR repairs 5-methycytosine deamination damage in a H3K36me3 dependent manner

The histone mark H3K36me3 is an important epigenetic modification involved in the recruitment of MMR to chromatin^36^. One of the hallmarks of H3K36me3 dependent MMR is differential repair of exons and introns where exons show significantly less mutations than expected due to increased H3K36me3 and MMR activity compared with introns^37^. To examine if MMR might play a role in the repair of 5mC deamination damage, we compared the observed and expected CpG C>T mutation densities in exons and introns in MBD4 deficient cancers and compared this to MMRd (i.e. MSI-H), MSS, and POLE mutant cancers (**Fig 5A-D**). Due to the proximity of introns and exons in the genome, comparison of their mutation density automatically controls for transcriptional activity and replication timing, both of which are also correlated with H3K36me3^38^. Exons have more observed and expected number of CpG C>T mutations compared with introns due to their generally higher GC content and CpG methylation levels^39^. Meanwhile, compared to introns, MSS and POLE showed substantial and significant decrease in observed exonic CpG C>T compared to expected (37.3% and 31.2%, p < 0.0001, one sample t-test, **Fig 5 A-B**), while this decrease was substantially less in MSI cancers due to the loss of MMR (2.74%, **Fig 5C**). Surprisingly, substantial and significant decrease in the observed exonic mutation density compared with expected exonic mutation density was also observed in MBD4 mutants (21.6%, p < 0.0001, one sample t-test, **Fig 5D**). As MBD4 is responsible for the repair for 5mC deamination damage, the decrease in observed exonic mutations in MBD4 mutants suggests that MMR may also be playing a role in the repair of G:T mismatches.

**Fig 5.**
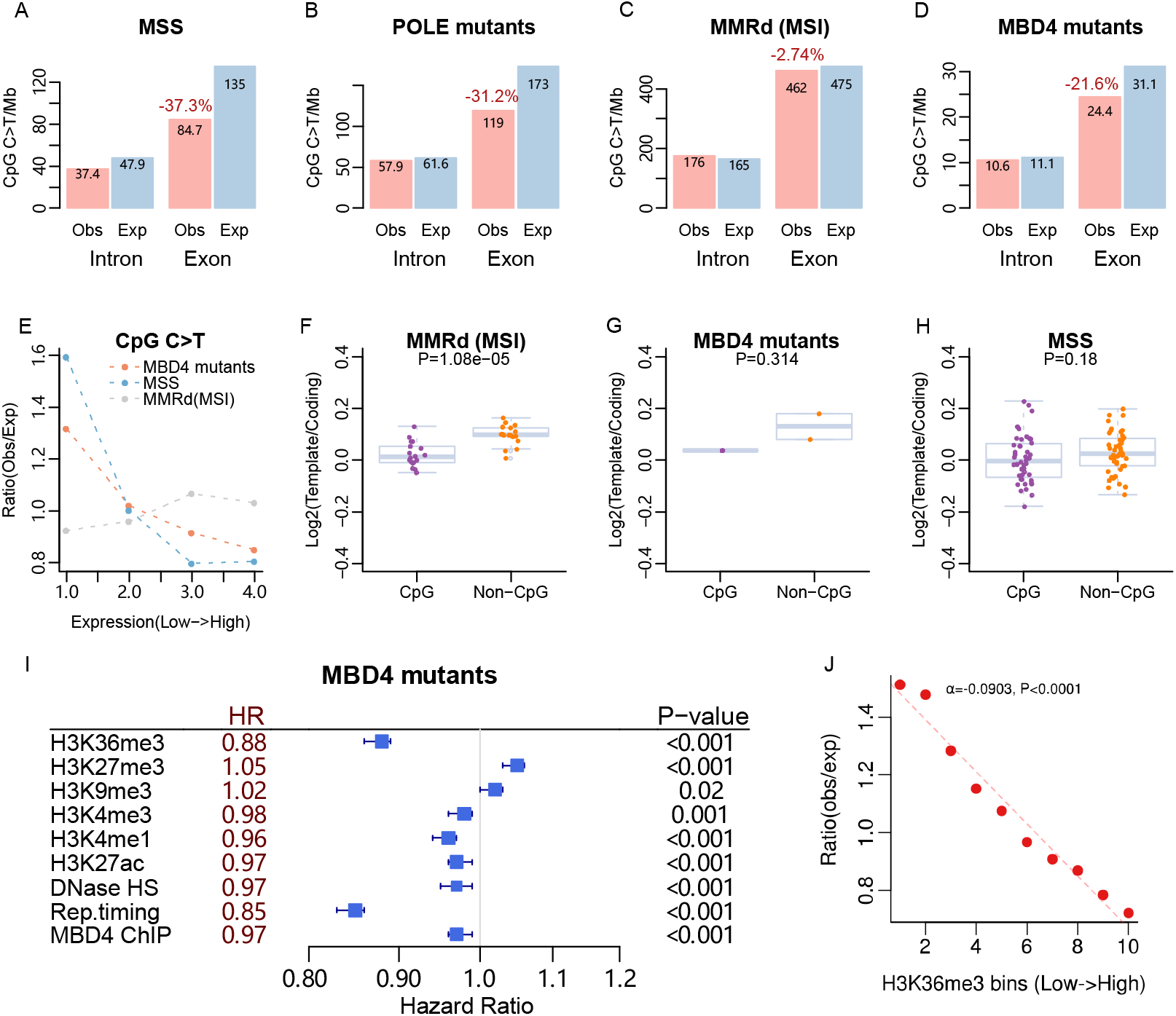
Association of mutation frequency with local determinants for different samples. **(A-D)** Observed and expected mutation densities in exons and introns across MSS, POLE mutants, MMR deficiency samples and MBD4 mutants. The expected mutations are obtained from simulation data that consider the abundance of tri-nucleotide mutational contexts. The decrease of observed and expected mutation density in exonic regions is indicated and calculated as (obs-exp)/exp. (**E**) Correlation of CpG C>T mutation ratio (obs/exp) with gene expression for MMR deficiency samples, MSS and MBD4 mutants. The P-values of the correlation are 7.7e-4, <2.2e-16 and 0.167 for MBD4 mutants, MMR deficiency samples and MSS respectively, and they were obtained from the linear regression model by fitting observed mutation density with unbinned gene expression. (**F-H**) Boxplot of transcription strand bias for CpG C>T and non-CpG C>T mutations in MMR deficiency samples, MSS samples and MBD4 mutants. (**I**) The hazard ratio of different epigenetics marks for CpG C>T mutation formation from multi-variable logistic regression model. 95% confidence level is indicated. P-value is calculated by Wald’s test. (**J**) Correlation between mutations in MBD4 mutants and H3K36me3 signal from mobilised CD34+ primary cells.

Recently, the MMR system has been reported to preferentially protect actively transcribed genes from mutation during transcription^13^. Consistent with this, we found that there was a negative correlation between CpG C>T mutation density and gene expression level for MSS and MBD4 mutants while there were more mutations in highly expressed genes for MMRd samples (**Fig 5E**). To determine if transcription coupled nucleotide excision repair (TC-NER) may also have a role in repairing 5mC deamination damage, we examined the transcription strand bias in MSI, MSS and MBD4 mutant cancers. We found that the transcription strand bias of CpG C>T mutations was close to zero in all cases (**Fig 5F-H**) suggesting that TC-NER is not involved in its repair. In MBD4 mutant cancers, this lack of transcription strand bias, further suggests that MMR is likely to be playing a role in the repair of the G:T mismatches. We next examined the association of CpG C>T mutation and different epigenetics marks for MBD4 mutants by multivariable logistic regression. Since CpG C>T mutations are highly dependent on methylation (**Extended Data Fig 6A**), we only selected mutations with highly methylated CpG (> 0.9) to ensure that we delineate the impact of histone modifications from DNA methylation. Apart from replication timing, we identified histone mark H3K36me3 had the lowest hazard ratio (HR) for CpG C>T mutation formation (HR = 0.88, p< 0.001, **Fig 5I, Supp Table 2**). There were fewer mutations in the regions of high H3K36me3 signal suggesting that in the absence of MBD4, the repair of CpG C>T mutations is dependent on H3K36me3 (**Fig 5J**). Interestingly, H3K36me3 also positively correlates with methylation level (**Extended Data Fig 6B)**. This suggests that in MBD4 mutants, although CpG methylation is the strongest determinant of mutation density, H3K36me3 activity is also an important factor for accounting for CpG mutation density. Taken together, these results suggest that even in the absence of MBD4, MMR has some capacity to facilitate the repair of 5mC deamination damage in a H3K36me3 dependent manner.

### MMR rather than MBD4 is essential for the repair of 5-methylcytosine deamination damage

While purified MBD4 can excise mismatched bases from DNA *in vitro*^40^, it is unclear whether MBD4 can repair 5mC deamination damage in the absence of MMR proteins *in vivo*. Previous studies have shown that MBD4 binding in the genome is strongly associated with DNA methylation density but is only weakly associated with H3K36me3^41^. To determine if MBD4 is able to facilitate the repair of 5mC deamination damage, we identified regions of the genome which are enriched in MBD4 and H3K36me3 based on ChIP-seq data from ENCODE. As the MBD4 ChIP-seq dataset is only available for the HepG2 cell line, we also used H3K36me3 ChIP-seq data from HepG2 to avoid cell type specific bias. After removing regions of low mappability, we identified 119,237 1kb windows in the genome that have either high (top 20%) MBD4 or H3K36me3 (**Fig 6A**). We also identified windows with low MBD4 or low H3K36me3. Using the respective regions, we compared the observed/expected mutation density across the windows in MBD4 mutants, MSS and MMRd (MSI) cancers. In high MBD4 regions, the observed/expected mutation rate was generally lower than the other regions (**Fig 6B**). This is due to the lower methylation level of CpGs in these regions despite overall high density of CpG, as well as the enrichment in early replicating regions (**Extended Data Fig 7A**). Importantly, the observed/expected mutation load was broadly similar across the three cancer types suggesting that MBD4 alone has little impact of observed mutation density. By contrast, for high H3K36me3 regions, in line with the importance of H3K36me3 for MMR, MSI had significantly more observed/expected mutations than MBD4 mutants and MSS cancers. The over 38% decrease in observed/expected mutation load in MBD4 reinforces our earlier results that MMR can repair 5mC deamination damage in the absence of MBD4. To further quantify the impact of MBD4 binding on 5mC deamination damage, we generated multivariable regression models and found that in MMRd cancers, MBD4 signal was not associated with lower likelihood of mutation (HR = 1.06, Figure 6C, see MutLα and MutSα separately in **Extended Data Fig 7B-C**). In MSS cancers, although a small decrease in HR was present for MBD4, H3K36me3 was a larger contributor to mutation likelihood (HR = 0.89 versus HR = 0.96, **Figure 6D**).

**Fig 6.**
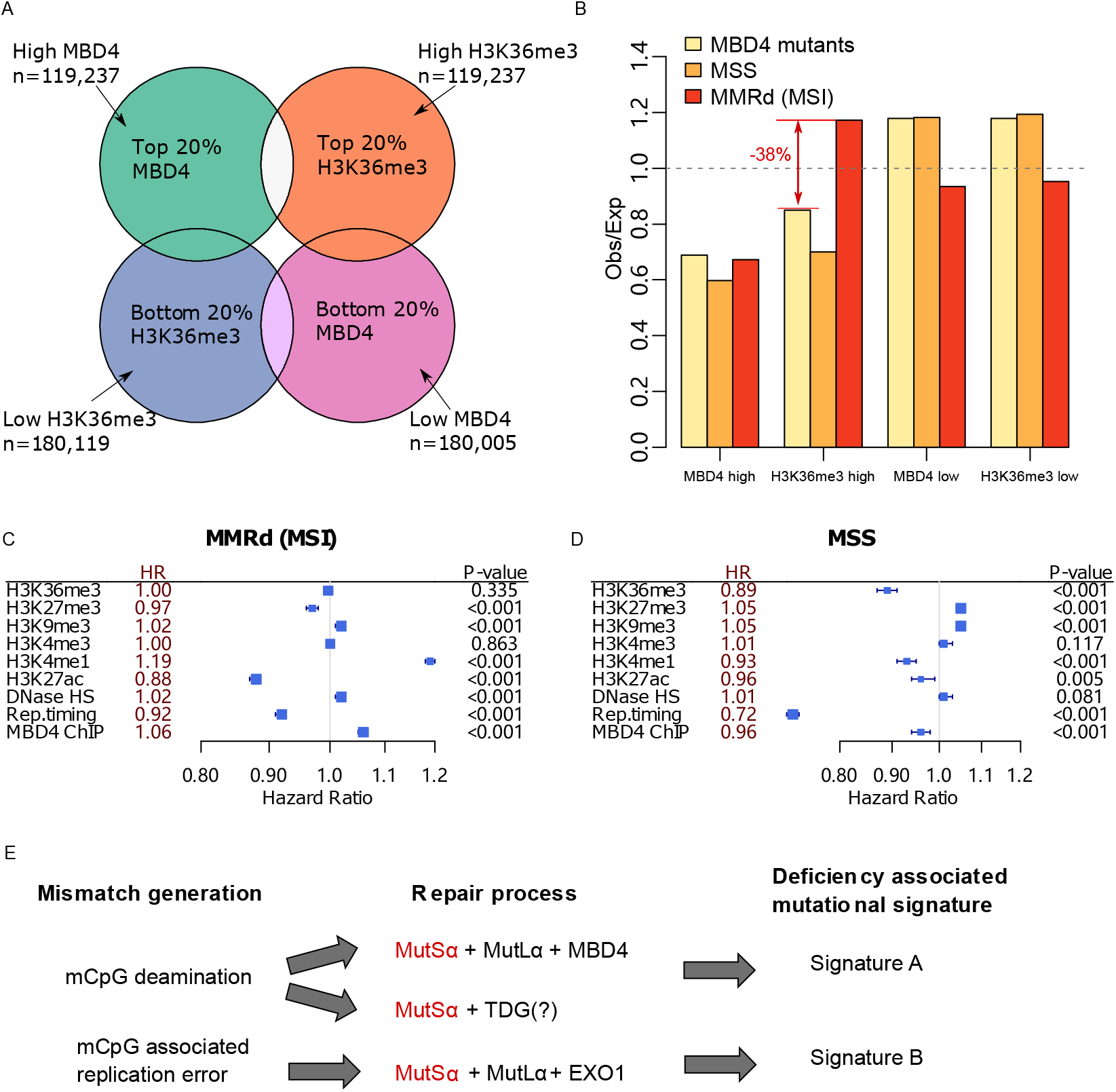
Association of MBD4 binding sites, histone mark H3K36me3 and mutations. (**A)** Venn diagram indicating the number of regions classified as top and bottom MBD4 and H3K36me3 signal based on the HepG2 cell line. (**B**) The ratio of observed and expected CpG C>T mutations in different regions for MBD4 mutants, MSS and MMRd cancers. The hazard ratio of different epigenetics marks for CpG C>T mutation formation from multivariable logistic regression model for MMRd (**C**) and MSS (**D**) cancers. 95% confidence level is indicated. P-value was calculated by Wald’s test. (**E**) Schematic of proposed mechanism of mismatch formation, repair and mutations.

It has been observed that truncating mutations in MBD4 are common in MSI cancers and truncated MBD4 can exert a dominant negative effect^42^. We have previously shown that MSI cancers with and without MBD4 truncating mutations does not show any difference in C>T mutation density at methylation CpG sites^35^. To examine this further in the context of MutLα and MutSα deficient MSI cancers, we found that 13.7% (27/197) of MutLα and 11.1% (2/18) of MutSα deficient cancers harboured MBD4 truncating mutations (**Extended Data Fig 7D, Supp Table3**). We did not find any difference in the mutational signature contributions in MBD4 wild-type and mutants for both MutLα and MutSα deficient cancers (**Extended Data Fig 7E**). We also examined the expression of MBD4 in TCGA MSI and MSS samples. While MBD4 expression was significantly lower in MutLα deficient cancers compared with MSS cancers (**Extended Data Fig 7F**), no association between MBD4 expression and SigA contribution was observed for MutLα deficient cancers (**Extended Data Fig 7G**). This further suggests that MBD4 is not a major factor in limiting CpG C>T mutations in MMRd cancers.

As MutSα deficient cancers have the highest SigA contribution and is responsible for the recognition of mismatched bases in DNA, our data suggests that it is the essential component for the repair of 5mC deamination damage. Interestingly, we found TDG to be upregulated in MutLα deficient cancers (**Extended Data Fig 7H**). This upregulation suggests that TDG may be the alternative glycosylase that can collaborate with MMR in the repair of 5mC deamination damage independently of MutLα and MBD4 (**Figure 6E**).

### Somatic mutations in MutSα mutant cell lines are largely caused by DNA replication errors

As MutSα deficient human cancers are highly enriched in SigA with a high proportion of CpG C>T mutations, we were intrigued that clonal mutations derived from two independent cultured MutSα mutant cell lines^43,44^ had a broader distribution of mutation types (**Fig 7A-B**), with both presenting a high contribution of SigB (**Fig 7C**). One is based on the human HAP1 cell line with knockout of MSH6 mediated by CRISPR-Cas9^43^ and the other one is based on human DLD-1 colon adenocarcinoma cells with MSH6 deficiency^44^. In both cases, somatic mutations were acquired in culture following clonal isolation and expansion. To determine if the CpG C>T mutations were likely to have formed in a similar way as in human cancers, the replication strand bias of CpG C>T and non-CpG C>T mutations for these cell lines was examined. Interestingly, unlike the human MMRd cancers, both MutSα deficient cell lines showed strong replication strand bias with no significant difference between CpG and non-CpG C>T mutations (p >0.05, Student’s t-test, **Fig 7D-E**). As cell lines replicate almost continuously and the mutations are acquired over the course of just one month, the result is consistent with the CpG C>T mutations forming during DNA replication. In contrast, cancer samples or pre-malignant cells replicate much more slowly than cell lines with mutations accumulating over many years, consistent with most CpG C>T mutations occurring via the deamination of 5mC, independent of DNA replication.

**Fig 7.**
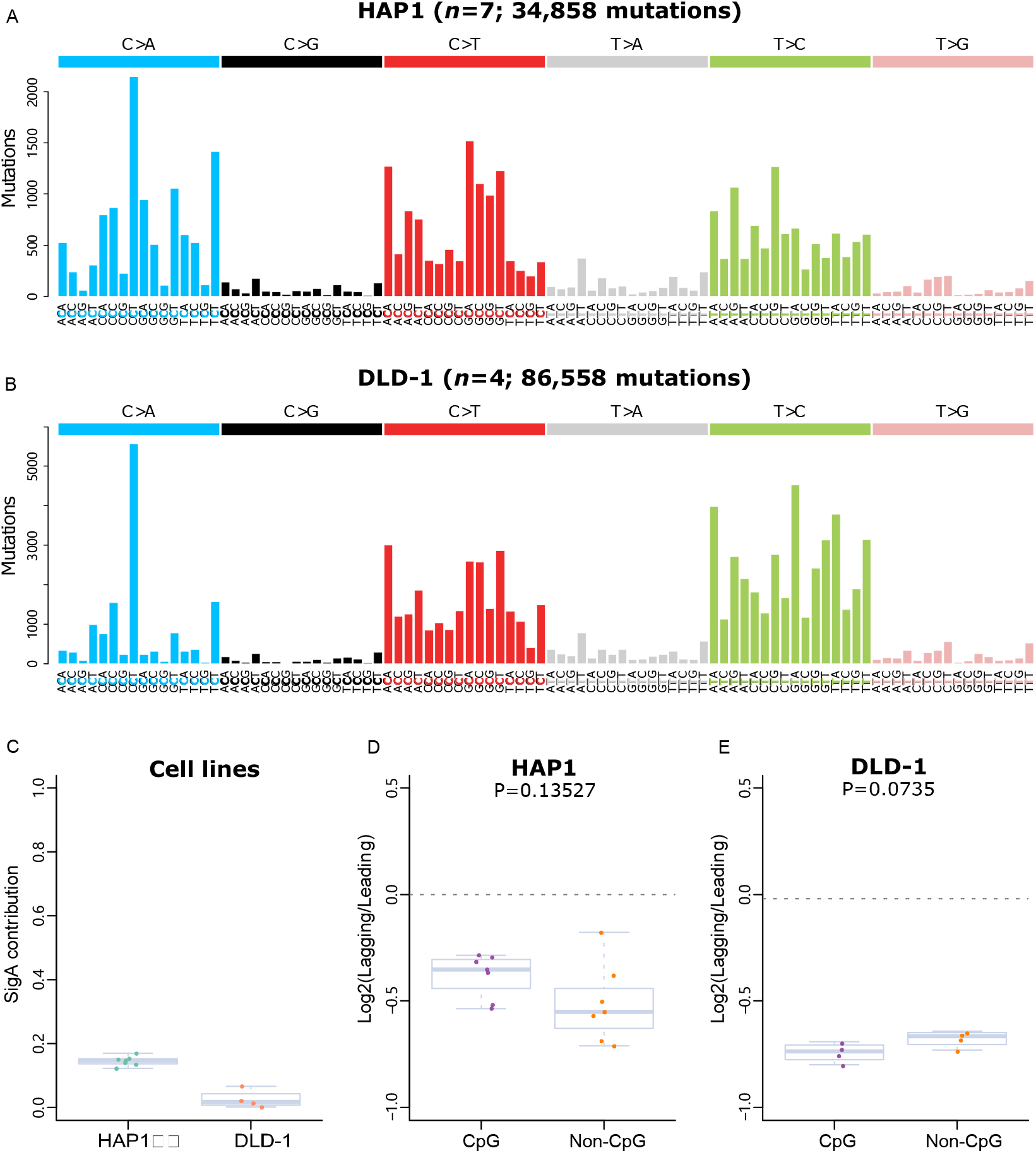
Mutation spectrum and replication asymmetry for MutSα mutant cell lines. Mutation spectrum for cultured HAP1 (A) and DLD-1 (B) cell lines. (C) SigA contribution for HAP1 and DLD-1 cell lines. Boxplot of replication stand bias for CpG C>T and non-CpG C>T mutations in HAP1 (D) and DLD-1 (E) cell lines.

## Discussion

A number of previous studies have sought to use genomics to determine the mechanisms underlying the different mutational signatures in MMRd cancers^4–6, 16, 45^. However, these efforts have been either restricted by cell line/organoid models^4,5^, non-mammalian models^45^, a lack of samples^6^ or incomplete assessment of all the data types available^16^. In this study, we carefully determined the status of the canonical MMR genes and classified each sample as being MutSα or MutLα deficient. In doing so, we found significant differences in the mutational signatures of MutSα and MutLα across four independent cohorts. Specifically, MutSα deficiency presents a high CpG C>T mutation spectrum, while MutLα mutants have a broader mutational spectrum including C>A, C>T and T>C mutations. These results are consistent with a recent published study of whole exome sequenced MSH2/MSH6-deficient gliomas which were all found to have a high frequency of CpG C>T mutation similar to SigA^46^. Another study based on the MSH6 null mouse model also reported elevated mutation frequency and predominance of G:C to A:T transition in MSH6 deficient small intestinal epithelium^47^.

Due to the susceptibility of cytosine (at CpG sites) to a variety of chemical modifications, CpG C>T mutations are common in cancer genomes and are generally recognised to be the result of 5mC deamination. Nevertheless, 5mC can also lead to CpG mutations in other ways^30,35,48^. While MMR is known to correct G:T and other mismatches that result from DNA replication errors, the repair of 5mC deamination should require excision of thymine from the G:T mismatch to restore the correct G:C pair^31^. It has been well established that MBD4 plays a critical role in the repair of mutations caused by 5mC deamination^32^. C>T mutations that arise from unrepaired 5mC deamination induced G:T should show no replication strand asymmetry as the deamination process should be largely independent of DNA replication (**Fig 4C**). By contrast, CpG C>T mutations that arise as a result of DNA replication errors, such as those in POLE mutant cancers^34^, display a high degree of asymmetry (**Fig 4D**). As the CpG C>T mutations observed in MMRd cancers, particularly those deficient of MutSα, lack strong replication strand asymmetry, this suggests that most of these mutations likely arise from replication independent 5mC deamination. As further evidence, we found that CpG C>T mutations generated by clonal expansion of MMRd cell lines show significant replication strand bias (**Fig 7D and E**), reflecting the rapid rate of cell division and lack of time for 5mC deamination induced mutations to accumulate. Thus, we propose that the two main mutational processes operational in MMRd cancers are a replication independent 5mC deamination induced C>T mutational signature (i.e. SigA) and a replication dependent DNA polymerase error driven signature (i.e. SigB) (**Fig 6E**). As SigA contributes to over 50% of mutations observed in most MMRd cancers (**Fig 2 C-F**), the inability to repair 5mC deamination damage can be considered the major mutational process in MMRd cancers.

Activities of MMR outside the context of DNA replication have been termed ncMMR and have been generally associated with the (error-prone) repair of DNA lesions through the recruitment of error-prone DNA polymerases^10–12^. It is therefore surprising that a form of ncMMR appears to participate in the error-free repair of 5mC deamination damage. Nevertheless, a recent study showed evidence of ncMMR dependent error-free repair of oxidative damage in actively transcribed genes^13^. Our data suggests that for the repair of 5mC deamination damage MMR is only performing the function of mismatch/lesion recognition to promote recruitment of other DNA repair factors pathways. Interaction of MMR and base excision repair has been studied *in vitro* in the context of active DNA demethylation^14^ and their results support a model where the two pathways collaborate to facilitate error-free removal of 5mC induced lesions including G:T mismatches in a MSH2 and TDG dependent manner. Our findings support this model. MutSα is known to have the role in DNA mismatch recognition and its MSH6 subunit contains the PWWP domain that binds H3K36me3 to facilitate its recruitment^36^. The high contribution of SigA for MutSα deficient cancers suggests that MutSα is essential for efficient mismatch recognition to recruit the MutLα-MBD4 complex, and potentially TDG, for the repair of 5mC deamination damage. Future studies will be required to fully elucidate the mechanisms of this form of ncMMR in *in vivo*.

In summary, we demonstrate that replication independent 5mC deamination contributes to most CpG C>T mutations in MMRd cancers. We find that MMR is in fact essential for the repair of 5mC deamination induced G:T mismatches. This non-canonical MMR function is likely to be MutSα dependent as MutSα deficient cancers are highly enriched in CpG C>T mutations. These results provide new insights of mutational processes in MMRd cancers and further our understanding of the ever-important MMR pathway.

## Acknowledgments

The authors would like to thank Ian Majewski and Rebecca Poulos for valuable feedback on the manuscript. This research is funded by Seed Funding from the University Grants Council, The University of Hong Kong to JWHW.

## Author Contributions

J.W.H.W conceived and designed the research; H.F, J.O. and J.W.H.W developed methodology and performed research; H.F, X.Z., J.O. J.A.B. and J.W.H.W analysed data; H.F. and J.W.H.W wrote the manuscript.

## Methods

### Data collection

A list of 316 MSI-H cancer samples including colorectal cancer (CRC), stomach adenocarcinoma (STAD) and uterine corpus endometrial carcinoma (UCEC) were obtained from TCGA Pan-Cancer dataset^49^. Other three independent cohorts were obtained for validation: Depmap cohort comprises 99 MSI-H samples across 16 cancer types^24^. MSK-CRC and MSK-UCEC cohorts contain 99 MSI-H colorectal cancers and 22 MSI-H endometrial cancers respectively, both with targeted sequencing data^22,23^. All these data are summarised in **Supp Table 4.**

### MutSα and MutLα classification

For the 316 samples from TCGA Pan-Cancer cohort, signature contributions were assigned by fitting COSMIC mutation signature v3 via the Sigfit R package^21^. Samples with high contributions (>10%) of signature SBS10a/b, SBS14 and SBS20 were excluded to avoid the effect of mutational processes that are caused by POLE and POLD1. DNA mutation, RNA expression and methylation data were applied to the remaining 266 samples for classification by the steps below: (1) Linear regression analysis was performed based on MLH1 methylation and expression. The regression equation was obtained as: y=9.050-4.996*x. Hypermethylated MLH1 is defined as β>0.25 and the low MLH1 expression cutoff value was obtained as 7.8 based on the equation. (2) Then MutSα and MutLα were determined based on the RNA expression and truncating mutation of the MMR genes that are elaborated in **Extended Data Fig 1.** For 99 MSI-H samples from the Depmap cohort, we first classified samples with truncating mutations in MSH2/MSH6 as MutSα if they have no aberration in MLH1/PMS2. Then the remaining samples with no aberration in MSH2/MSH6 were classified as MutLα (**Supp Table 5**). Due to the availability of data for MSK-CRC cohort, the classification of MutSα and MutLα is based on DNA mutations of MMR genes. Samples with truncating mutations in MSH2/MSH6 and without truncating mutations in MLH1/PMS2 are classified as MutSα. The remaining samples are classified as MutLα (**Supp Table 5**6. For samples from the MSK-UCEC cohort the classification is based on immunohistochemistry of the four MMR genes (**Supp Table 7**).

### Mutation simulation at tri-nucleotide resolution

Mutation simulation was performed by SigProfilerSimulator^50^ to preclude the bias of tri-nucleotide composition which could affect the mutation distribution in local regions. Briefly, the total number of simulated mutations for each sample is equal to the observed mutations, but the position of the mutation is relocated according to the frequency of tri-nucleotide context along the given region. Each sample was simulated 100 times and all the mutations are combined as expected mutations for subsequent local mutation density analysis.

### Mutational signature analysis

The profile of each signature was represented by six substitution subtypes: C>A, C>G, C>T, T>A, T>C and T>G. For signatures generated by tri-nucleotide context, each substitution on the cancer genome was examined by incorporating information on the bases immediately 5’ and 3’ to each mutated base to generate 96 possible mutational types. *De novo* signatures were extracted by Sigfit which applies a Bayesian inference algorithm ^21^. Mutational signatures were displayed and reported based on the observed tri-nucleotide frequency of the human genome. For validation cohorts, contribution of *de novo* signatures for each sample was calculated by fitting the mutations to the extracted *de novo* signatures.

### Replication timing and mutation density

The replication time of different chromosome regions was obtained for the HepG2 cell line from the ENCODE data portal^51^. Exome sequence with known replication time was divided into five bins from late to early: [-3.88, 51.98), [51.98, 66.30), [66.30, 74.95), [74.95, 80.74), [80.74, 87.95]. The counts of mutations within each bin were calculated as observed mutation. Similarly, the expected mutation counts were also computed for each bin based on simulated data. The slope was obtained from the linear regression model based on the correlation of mutation ratio (obs/exp) and replication timing.

### Gene expression and mutation density

The general gene expression data were obtained from GTEx Portal and all expressed genes were integrated into four bins according to the expression quartile. For each bin, only sites located within early replicated regions are adopted. The size of each bin was calculated based on the length of exons of each gene. The count of observed and expected mutations was calculated for each bin to determine mutation density.

### Calculating strand asymmetries

Replication direction was defined using replication timing profiles from a previously published paper^52^. Left (leading)- and right (lagging)-replicating regions were determined by the derivative of the profile. For a given mutation type in the right replication direction, the mutation counts (*N_1_*) in that region were calculated, and its complementary mutation was calculated as *n_1_*. Correspondingly, the mutation counts of this mutation type in left replication direction were calculated as *N_2_*, and its complementary mutation was calculated as *n_2_*. Then, asymmetry (*A*) was calculated as:

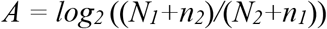

For the transcription strand asymmetry, coding and template strand were obtained from a published study^53^ and the asymmetry is reported as log2 ratio of (mutation count within template regions) / (mutation counts within coding regions).

### Computing mutation density in exonic and intronic regions

All gene coordinates were obtained from the UCSC table browser. Middle exons and middle introns were extracted for each gene. Then, the mutation density was calculated as mutation counts per megabase for both exonic and intronic regions.

### Associations between MBD4 mutant mutation density, histone marks and CpG methylation

As MBD4 mutants are derived from acute myelocytic leukemia, we obtained whole genome bisulfite sequencing data for E050 (Mobilized CD34 Primary Cells). Histone marks including H3K36me3, H3K27me3, H3K9me3, H3K4me3, H3K4me1, H3K27ac as well as DNase I hypersensitive site are derived from common myeloid progenitor and CD34-positive samples. For the data to estimate regression model for MSS and MMRd (MSI) samples, the mutations are from TCGA CRC cancer. Histone marks as well as DNase I hypersensitive site are obtained from Homo sapiens large intestine male embryo (108 days). All these data are downloaded from the Roadmap Epigenomics Atlas^54^. MBD4 ChIP-seq data obtained from HepG2 cells from ENCODE^51^. Only sites with methylation value >0.9 are adopted for fitting the logistic model. For the correlation of CpG methylation, MBD4 mutants mutation and H3K36me3 signal, all cytosines in the CpG di-nucleotide were merged into 12 bins according to their methylation level as: [0], (0, 0.1],…, (0.9, 1.0), [1]. These bins were then used as intersected regions to calculate mutation density in each methylation level. H3K36me3 bins were set based on the H3K36me3 signal. For the grouping of MBD4 and H3K36me occupied regions, the H3K36me data were also obtained from HepG2 from ENCODE^51^. MBD4 and H3K36me3 signal/input were calculated across 1kb windows across the genome. Regions that had an average mappability score of <1 based on UCSC *Duke Uniquness* 35 bp and those that overlapped *DAC blacklisted regions* were removed from the analysis.

## Supplementary tables

Supp Table 1. Sample information for TCGA cohort.

Supp Table 2. Multivariable logistic regression model.

Supp Table 3. MBD4 truncating mutation annotation in MSI samples.

Supp Table 4. Data cohorts summary.

Supp Table 5. Sample information for Depmap cohort.

Supp Table 6. Sample information for MSK-CRC cohort.

Supp Table 7. Sample information for MSK-UCEC cohort.

