## Supplementary Figures for "Deficiency in DNA mismatch repair of methylation damage is a major mutational process in cancer"

### Extended Data Figure 1

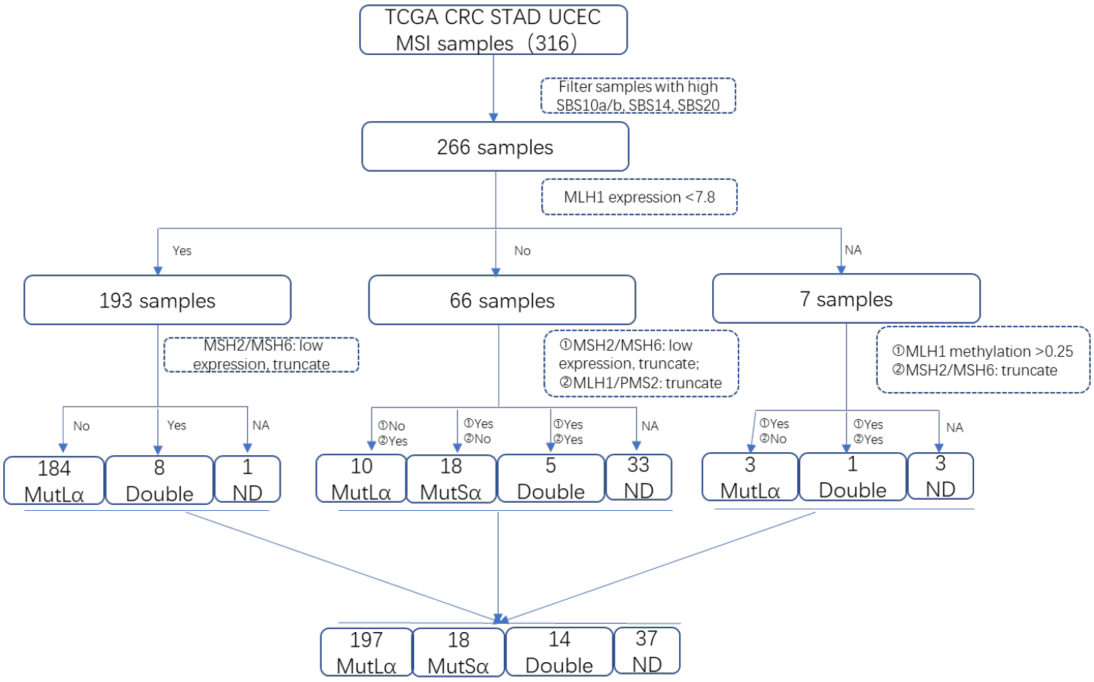


**Extended Data Fig 1. Diagram of MutSα and MutLα classification for TCGA MSI-H cancer samples.**

**Extended Data Figure 2**

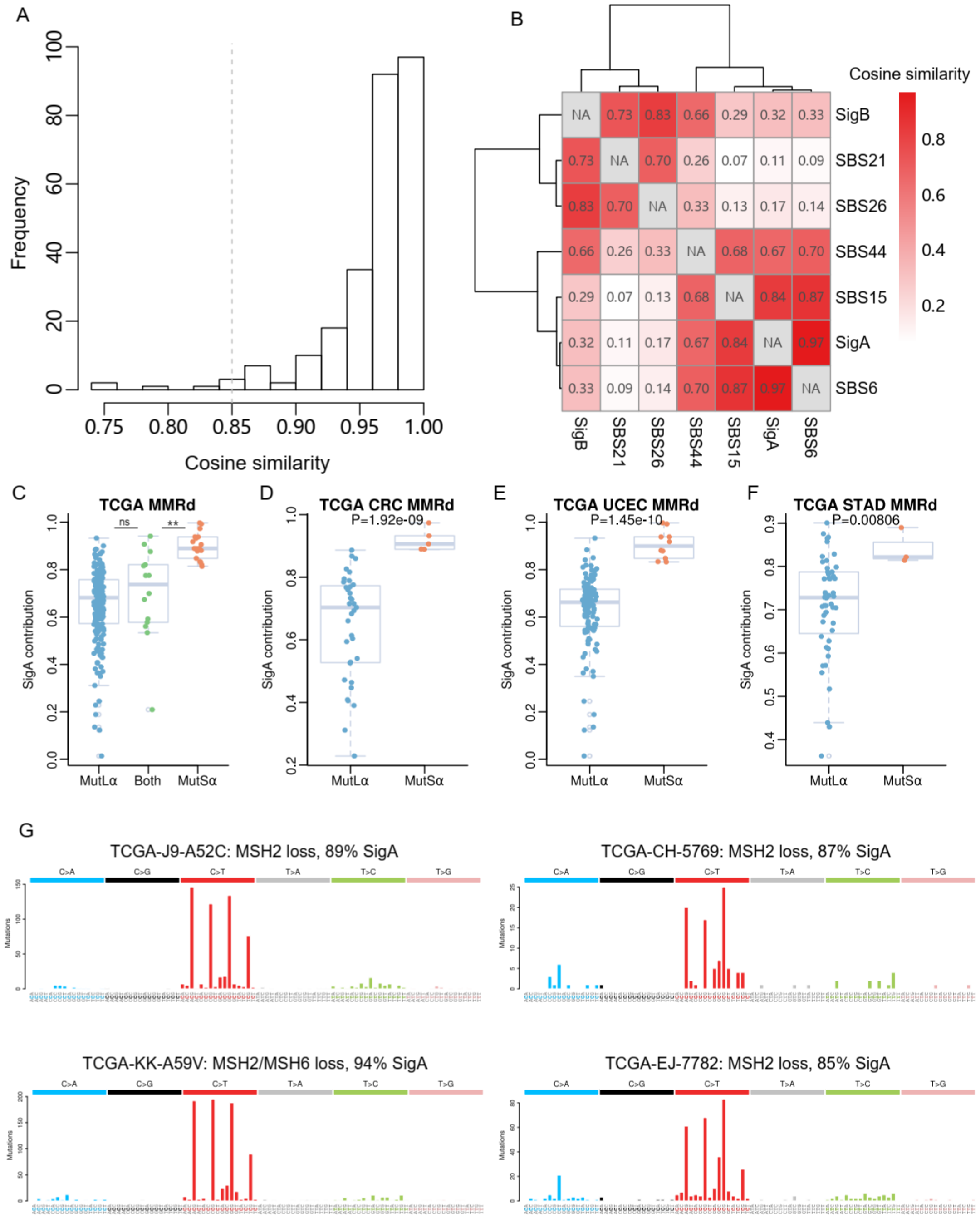


**Extended Data Fig 2. Characteristic of extracted de novo signatures and their contribution to MutSα and MutLα across different cancer types.** (**A**) Histogram of cosine similarity across MMR deficiency samples. Cosine similarity is calculated from the comparison of real mutational spectrum with reconstructed spectrum. (**B**) Hierarchical clustered heatmap of cosine similarity of two *de novo* signatures and five mismatch repair deficiency associated SBS signatures from COSMIC database. (**C**)

Boxplot of SigA contribution in MutSα, MutLα and Both mutants. (**D-F**) Boxplot of SigA contribution in MutSα and MutLα across three different cancer types including CRC, UCEC and STAD. (G) Mutation spectrum of four MutSα prostate cancer. P-value is calculated by two-tailed Student's t-test. ** $<0.01$, n.s. $>0.05$.

**Extended Data Figure 3**

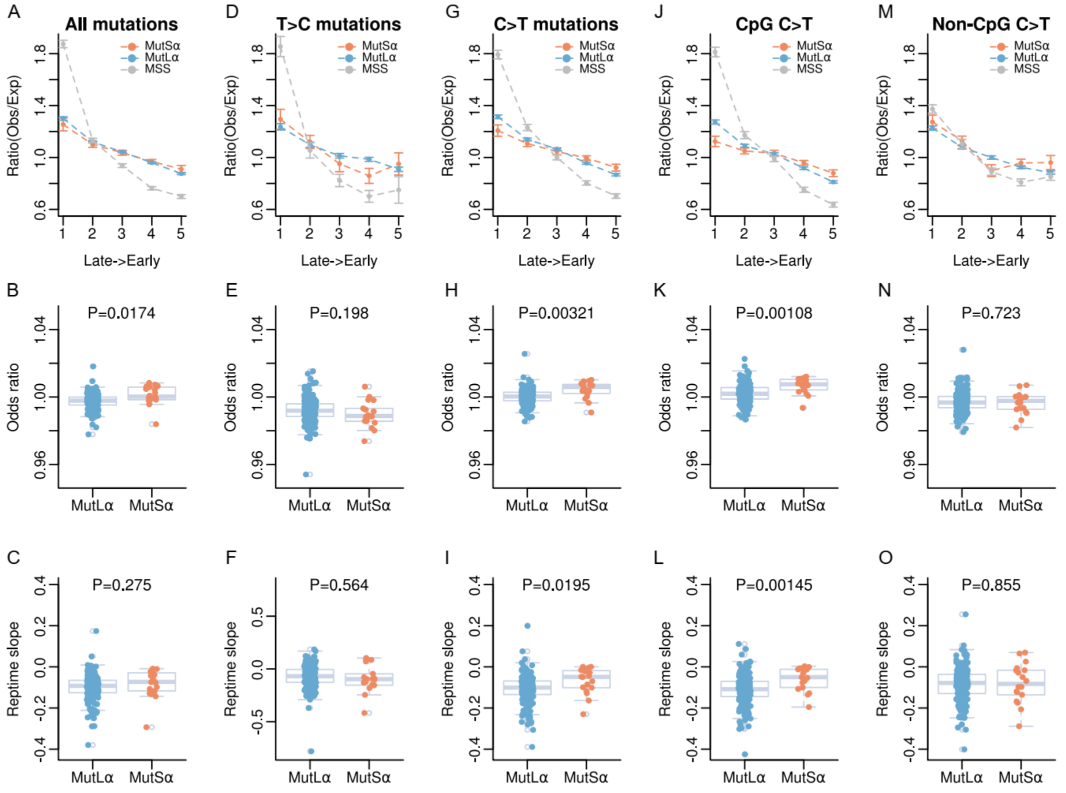


**Extended Data Fig 3. The association of mutation density with replication timing.** The association of total mutations (**A**), T>C mutations (**D**), C>T mutations (**G**), CpG C>T mutations (**J**) and non-CpG C>T mutations (**M**) with replication timing for MutSα, MutLα mutants and MSS samples. Error bars indicate +/- 2 SE. Boxplot of odds ratio for total mutations (**B**), T>C mutations (**E**), C>T mutations (**H**), CpG C>T mutations (**K**) and non-CpG C>T mutations (**N**) for individual sample in MutSα and MutLα mutants. Odds ratio is computed from logistic regression model that is estimated by mutations and replication timing for each sample. Boxplot of replication slope for total mutations (**C**), T>C mutations (**F**), C>T mutations (**I**), CpG C>T mutations (**L**) and non-CpG C>T mutations (**O**) for individual sample in MutSα and MutLα mutants. The replication slope is computed from the regression line by fitting mutation ratio (obs/exp) with replication timing bins. P-value is calculated by two-tailed Student's t-test.

**Extended Data Figure 4**

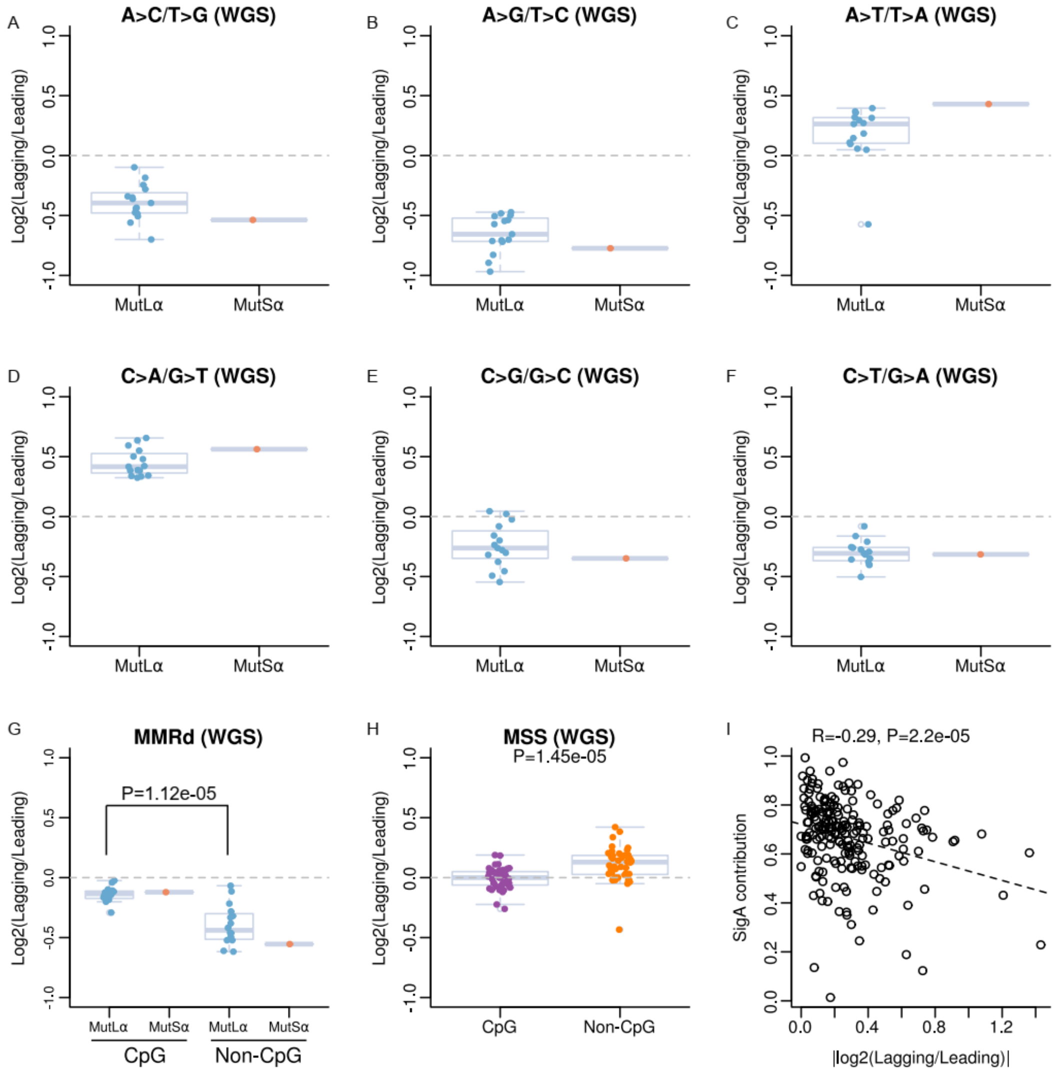


**Extended Data Fig 4. Replication asymmetry in MMRd and MSS WGS data and its correlation with SigA contribution. (A-F)** Replication stand bias of six types of mutations for MutLα and MutSα whole genome sequencing data. (**G-H**) Replication stand bias of CpG and non-CpG C>T mutations for MMRd and MSS whole genome sequencing data. (I) The correlation of SigA contribution and CpG C>T replication strand bias. The strand bias is calculated as an absolute value of the log2 ratio of mutation counts in lagging strand and leading strand.

**Extended Data Figure 5**

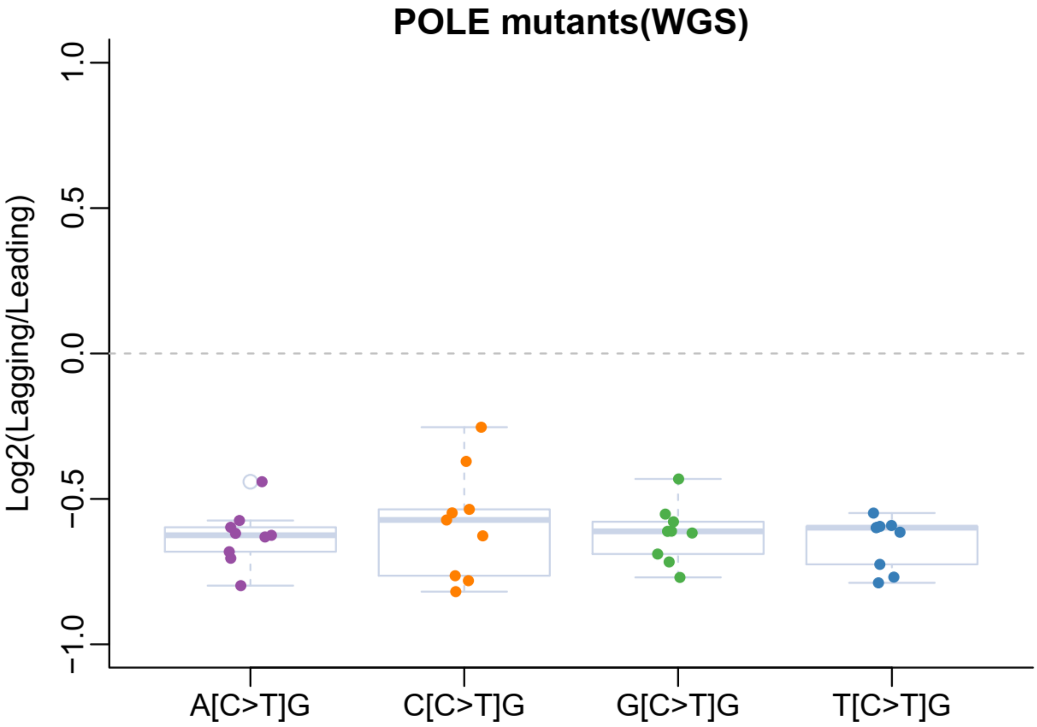


**Extended Data Fig 5. The strand bias of four CpG C>T trinucleotide contexts for POLE mutants.**

**Extended Data Figure 6**

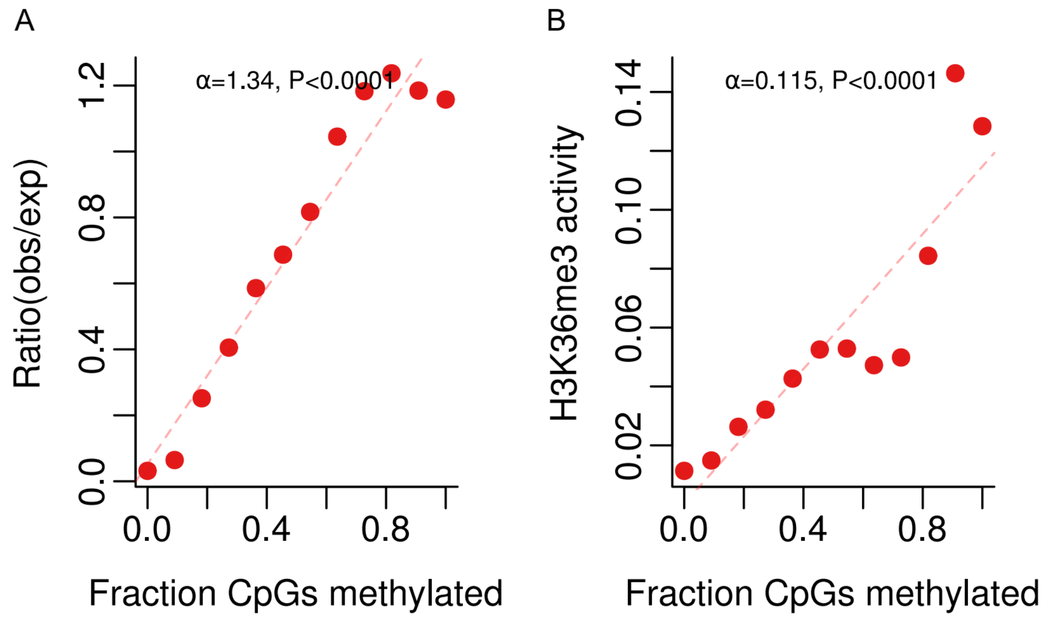


**Extended Data Fig 6. Correlation between mutation, CpG methylation and H3K36me3 signal in MBD4 mutants. (A)** Correlation between observe/expected mutations in MBD4 mutants and CpG methylation from mobilised CD34 Primary Cells. **(B)** Correlation between CpG methylation and H3K36me3 from mobilised CD34 primary cells.

**Extended Data Figure 7**

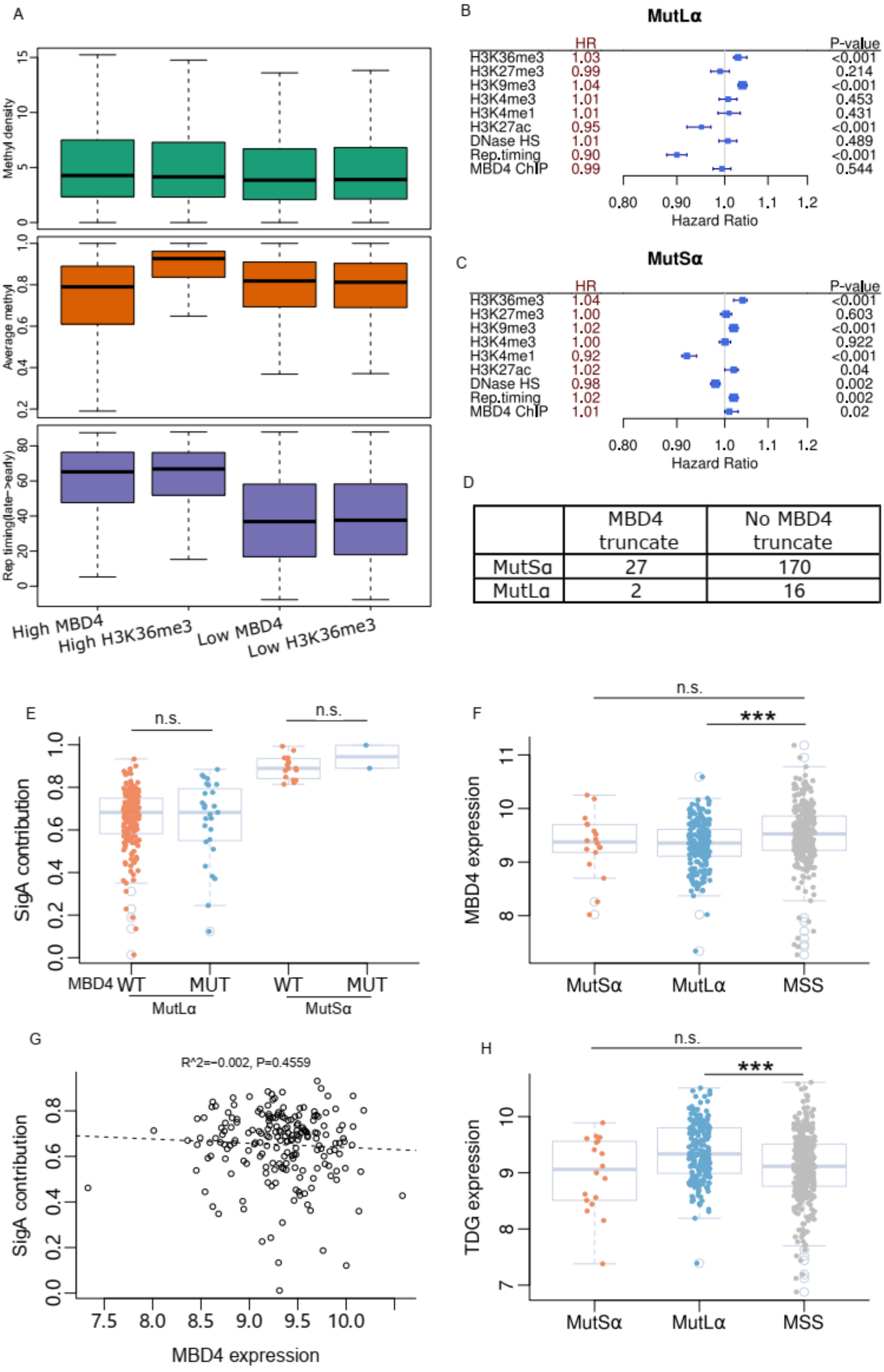


**Extended Data Fig 7. Contribution of MBD4 to mutation formation. (A)**

Methylation density, average methylation and replication timing distribution across High MBD4, High H3K36me3, Low MBD4 and Low H3K36me3 regions. **(B-C)** The hazard ratio of different epigenetics marks for CpG C>T mutation formation from multivariable logistic regression model for MutLα and MutSα cancers. 95% confidence level is indicated. P-value is calculated by Wald's test. (**D**) Contingency table of MBD4 truncated samples in MutLα and MutSα cancers. (**E**) SigA contribution in MBD4 WT (no truncation) and MBD4 MUT (truncating mutation) mutants for MutLα and MutSα cancers. (**F**) MBD4 expression in MutLα, MutSα and MSS cancers. (**G**) Correlation of SigA and MBD4 expression in MutLα cancers. (**H**) TDG expression in MutLα, MutSα and MSS cancers. P-value is calculated by two-tailed Student's t-test. ***<0.001, n.s. >0.05.
